# Thymic Stromal Lymphopoietin Promotes Proliferation and Contractility of Human Pulmonary Artery Smooth Muscle

**DOI:** 10.1101/441188

**Authors:** Michael Thompson, Venkatachalem Sathish, Logan Manlove, Benjamin Roos, Bowen Wang, Robert Vassallo, Jordan Miller, Christina M. Pabelick, Y.S. Prakash

**Author notes:** **Corresponding Author:** Y.S. Prakash, MD, PhD, Professor of Anesthesiology and Physiology, Department of Anesthesiology and Perioperative Medicine, 4-184 W Jos SMH, Mayo Clinic, 200 First St SW, Rochester, MN 55905.

## Abstract

Hypoxia is a well-recognized risk factor in several pulmonary vascular diseases including pulmonary hypertension (PH). Furthermore, hypoxia-associated inflammatory changes enhance the structural and functional changes in the pulmonary artery (PA) of PH patients. Understanding the mechanisms that link hypoxia and inflammation, particularly early in disease, is key to development of novel therapeutic avenues for PH. Thymic stromal lymphopoietin (TSLP) is an “early” inflammatory mediator thought to be critical in diseases such as asthma, chronic obstructive pulmonary disease and atopic dermatitis. TSLP has canonical effects on the immune system, but can also have non-canonical effects on resident lung cells, e.g. airway smooth muscle. Currently, the expression and role of TSLP in the PA is unknown. We hypothesized that locally-produced TSLP potentiates the effects of hypoxia in PA remodeling and contractility relevant to PH. Experiments in human PA endothelial cells (PAECs) and smooth muscle cells (PASMCs) found PAECs to be a larger source of TSLP which targets PASMCs to enhance intracellular Ca^2+^ responses to vasoconstrictor agonist as well as cell proliferation, acting via a number of signaling cascades including Stat3 and PI3/Akt. Hypoxia, acting via HIF1α, enhanced PAEC production of TSLP, and promoted TSLP effects on PASMCs. Interestingly, TSLP per se enhance HIF1a. Overall, these novel data highlight a role for TSLP in hypoxia effects on the PA, and thus relevance for inflammation in PH.

## Introduction

Thymic stromal lymphopoietin (TSLP) is an IL-7-like cytokine first identified in the thymus as a factor in T and B cell development,[1–3] but now found to be produced by a variety of non-thymic cell types including epithelial cells of the lung, gut and skin, fibroblasts, and circulating and tissue immune cells, particularly dendritic cells.[2, 4, 5] Acting via a heterodimeric complex of its receptor TSLP-R and IL7Ra, TSLP serves as an interface between the environment and the body to skew the immune response towards a Th2 phenotype early during the response to allergic and other stimuli [6, 7]: an effect that has driven the high interest in TSLP in atopic dermatitis, allergic asthma, and non-infectious GI disorders.[3, 8–11] In spite of such interest, there are currently significant knowledge gaps in TSLP expression and signaling patterns (particularly given species and cell type differences).

The relevance of TSLP to the pulmonary vasculature lies in the role of inflammation in diseases such as pulmonary hypertension (PH). While different types of PH differ in etiology, risk factors and presentations, there are two factors that may be important across multiple PH groups: hypoxia and inflammation. Hypoxia is certainly a key aspect of Group III PH, and can also be contributory in Group IV.[12, 13] In addition, there is increasing evidence that inflammation also plays a significant role in PH pathogenesis and exacerbation, and is likely important not only in Group III, but also Group I and IV.[14, 15] In this regard, hypoxia can influence the inflammatory response with subsequent effects on PA structure and function.[16–18] For example, in high-altitude mountain sickness,[19, 20] circulating levels of pro-inflammatory cytokines are increased. In healthy persons, hypoxia increases serum IL-6, and C-reactive protein as well IL-6 receptor levels.[21–23] In mice, even short-term hypoxia leads to vascular leakage, inflammatory cell infiltrates and elevated serum cytokines.[24, 25] There is strong circumstantial evidence for an inflammatory pathogenesis of PH[24, 26–28]. Furthermore, PH is associated with inflammatory conditions including rheumatoid arthritis, lupus, and collagen vascular diseases.[24, 29, 30]

The significance of understanding the mechanisms by which hypoxia induces inflammation is clear. While such mechanisms may vary in different forms of PH, what is important to recognize is that if we can identify early mediators of the inflammatory response to hypoxia, then novel preventive and therapeutic avenues can be explored. Here, given the increasing recognition that TSLP is an early respondent and inflammatory mediator in other organ systems, we believe that the TSLP/TSLP-R signaling cascade is a novel target to explore in PH, especially given the potential that TSLP may have pleiotropic effects relevant to PH pathophysiology. Furthermore, the information derived from our studies has the potential to target other disease conditions where the hypoxia-inflammation axis is important, including lung injury especially following transplantation. Accordingly, in the current study, we explored the potential role of TSLP in the human pulmonary artery (PA) as a first step towards understanding the contribution of TSLP to PH. We hypothesized that TSLP represents a locally-produced inflammatory mediator in the PA, with autocrine/paracrine effects on the endothelium and smooth muscle.

## Materials and Methods

### Culture of Human Pulmonary Artery Endothelial Cells (PAECs) and Smooth Muscle Cells (PASMCs)

Human PAECs and PASMCs were obtained from commercial sources (Thermo Fisher Scientific, Waltham, MA and ATCC, Manassas, VA) or, under a Mayo Institutional Board-approved protocol conforming with the Declaration of Helsinki. Cells were isolated from PA of lung samples incidental to patient thoracic surgery at Mayo Clinic-Rochester (lobectomies, pneumectomies for non-PH transplant indications and focal non-infectious indications such as localized tumors) as previously described [31–33]. Since samples were obtained post-hoc incidental to surgery and not for the purpose of this research *per se,* and furthermore patient care was unaffected by any studies performed with such samples, the protocol was considered not Human Subjects research and exempt by Mayo IRB (minimal risk protocol).

Sample collection was limited to normal appearing areas of the vasculature identified by the pathologist and verified under gross microscopy. Samples were de-identified and were considered not Human Subjects Research (minimal risk protocol). The PA was transported rapidly to the laboratory in ice-cold Hank’s Balanced Salt Solution (HBSS), cleaned and the endothelium separated, with adventitia removed for further cell isolation.

For PASMCs, endothelium-denuded tissue was minced and placed in 100mm petri dishes with DMEM F12 containing 10% FBS/1% ABAM. Tissue explants were maintained for 5-7 days at 37°C in 95% air/5% CO_2_ after which the source tissue was removed, cells grown to confluence for experiments, plated in 60mm dishes (Western analysis), 8 well Lab-Tek culture chambers or 96 well plates (proliferation assays).

PAECs were isolated via modification of methods described previously.[31, 34] Briefly, endothelium from PA was minced with a razor blade and incubated in Earle’s Balanced Salt Solution (EBSS) containing 0.1% Collagenase II and 0.25 U/ml Dispase (Thermo) for 30 min at 37°C with continuous agitation. DNase I (7.5 μg/ml final concentration, Sigma, St. Louis, MO) was then added and tissue incubated for another 30 min at 37°C. Dissociated cells were separated from undigested tissue using a 100μm strainer and pelleted at 400*xg* for 5 minutes. Cells were resuspended in PBS and subsequently incubated with anti-CD45 and anti-CD31 microbeads (Miltenyi Biotec, Inc, Auburn, CA) and endothelial cells magnetically separated using the AutoMacs Pro cell separation system (Miltenyi) according to manufacturer’s protocol. Final cell pellet was resuspended in Endothelial Cell Growth Medium-2 (EGM-2, Lonza, Walkersville, MD), plated in 75cm^2^ flasks and grown to confluence.

Prior to experimentation, PAECs and PASMCs were serum-deprived for 24 h and all cells were used between passages 1 and 5. PAEC and PASMC phenotype was verified by immunostaining using anti-CD31 (Abcam, Cambridge, MA; ab24590) or smooth muscle specific anti-smooth muscle actin (Sigma, St. Louis, MO; A2547) primary antibodies, respectively.

### Cell Exposures

Human PAECs and PASMCs were incubated for 72 h at 37°C in normoxia (21% O_2_) or hypoxia (1% O2) in serum free medium (control), supplemented as appropriate with 20ng/ml recombinant human TSLP (R&D Systems, Minneapolis, MN). The role of HIF-1α was verified using 10μM (final concentration) of HIF-1 pharmacological inhibitor (Santa Cruz Biotechnology, Inc., Dallas, TX). The functional activity of TSLP was inhibited with the use of 10μM STAT-3 Inhibitor, LLL12 (EMD Millipore, Billerica, MA). Inhibitors were incubated with cells 1h prior to adding TSLP.

### Immunostaining

PAECs and PASMCs were seeded in 8 well Lab-Teks (Thermo) at 5,000 cells/well. Cells were fixed in 4% paraformaldehyde for 15 min and washed with TBS. For detection of TSLP cells were permeabilized for 10 min in TBS containing 0.1% Triton X-100. Detection of TSLP-R did not require permeabilization. Cells were then blocked for 1h in TBS containing 4% normal donkey serum and incubated for 1h at room temperature with 5μg/ml primary antibody (TSLP, Santa Cruz, sc-33791; TSLP-R, Santa Cruz, sc-83871). Samples were then washed with TBS and incubated with donkey anti-rabbit Alexa 555 secondary antibody (1:500, Thermo) for 1h at room temperature, washed with TBS and coverslips mounted with Flurogel II containing DAPI (Electron Microscope Sciences, Hatfield, PA). Cells were visualized using a Nikon Eclipse TE2000-U microscope with a 40x/1.30 NA oil objective lens.

### Assessment of TSLP Levels

Cell culture supernatants from PAECs and PASMCs grown on 60mm petri dishes and exposed to normoxia or hypoxia, were concentrated using Ultracel 3k Amicon Ultra centrifugal filters (EMD Millipore) and assayed for TSLP via ELISA (R and D Systems, Minneapolis, MN) according to manufacturer’s protocol, and as described previously for airway cells.[5] Changes in optical density were determined using a FlexStation3 microplate reader (Molecular Devices, Sunnyvale, CA) set to 450 nm (wavelength correction set to 540 nm) and compared to manufacturer-provided calibration curve.

### Western Analysis

Standard SDS-PAGE (Criterion Gel System; Bio-Rad, Hercules, CA; 4-15% gradient gels) and Trans-Blot Turbo transfer system using nitrocellulose membranes (Bio-Rad) were used. Membranes were blocked with LiCor blocking buffer (LiCor, Inc., Lincoln, NE) for 1h at room temperature prior to addition of primary antibody (1 μg/mL rabbit anti-TSLP-R, sc-83871,Santa Cruz; 1 μg/mL rabbit anti-TSLP, sc-33791, Santa Cruz;1 μg/mL rabbit anti-ERK, sc-94, Santa Cruz; rabbit anti-PI3K, 1:500, 06-195, EMD Millipore; 1μg/ml rabbit anti-Akt, , 4691, Cell Signaling Technologies, Inc, Danvers, MA; 2 μg/mL rabbit anti-Cyclin E, sc-25303, Santa Cruz; 2 μg/mL rabbit anti-PCNA, sc-25280, Santa Cruz; 2 μg/mL rabbit anti-JAK2, sc-294, Santa Cruz; 2 μg/mL rabbit anti-STAT3, sc-482, Santa Cruz ; 1μg/ml rabbit anti-GAPDH, 2218, Cell Signaling) overnight at 4°C with gentle rocking. Membranes were washed in TBS, incubated with goat anti-rabbit or anti-mouse secondary antibodies (IRdye800, 1:10,000 dilution, LiCor) for 1h at room temperature. Blots were visualized and densitometry performed with an Odyssey infrared imaging system (Li-Cor Biosciences).

### Cell Proliferation Assay

Proliferation of PASMCs was assayed at 72h (with or without preceding interventions) using the CyQuant NF kit (Invitrogen) according to manufacturer’s protocol. Cells were first washed with HBSS and exposed to the CyQuant dye for 1 h at room temperature. Dye binding to DNA (fluorescence) was measured on the FlexStation3 microplate reader. Dye calibrations were performed empirically using different cell counts to establish a standard curve and fluorescence converted to cell number to determine degree of proliferation.

### Real Time Calcium Imaging

We have previously described the techniques for real-time fluorescent imaging of [Ca^2+^]_i_ using 5 μM Fura-2/AM [31, 35]. Following dye loading for 45 minutes at room temperature, cells were visualized using a Nikon TE2000-U inverted microscope and Nikon Elements imaging software. Cells were initially perfused with HBSS containing 2mM CaCl_2_ to establish a baseline then perfused with 10μM serotonin. [Ca^2+^]_i_ responses from 20 regions of interest were obtained from multiple cells per chamber.

### Statistical Analysis

All experiments were performed in cells from 4-8 different artery (patient) samples (n values are indicated for each experiment in the results). Not all protocols were performed in each sample although a minimum of 4 different samples was used for each experiment. For box plots, both median (solid line) and mean (dashed line) are indicated. Analysis of results was accomplished using one-way ANOVA with repeated measures or Tukey post-hoc analysis where appropriate. Statistical significance was set at p<0.05; all values are expressed as means ± SE.

## Results

### TSLP and TSLP-R expression and secretion in human PA

Immunofluorescence staining of human PAECs and PASMCs for TSLP and TSLP-R showed the presence of TSLP and TSLP-R in both cell types (Figure 1A). Extracellular TSLP measured by ELISA demonstrated that PAECs release TSLP (110±24 pg/ml at normoxia), and such release was significantly enhanced (252±91 pg/ml) following 72h hypoxia (p<0.05 compared to normoxia control; n=8). However, hypoxia did not appear to influence PASMC TSLP release (68±8 and 80±15 pg/ml, respectively; n=5) (Figure 1B): levels that were lower than those shown by PAECs.

**Figure 1.**
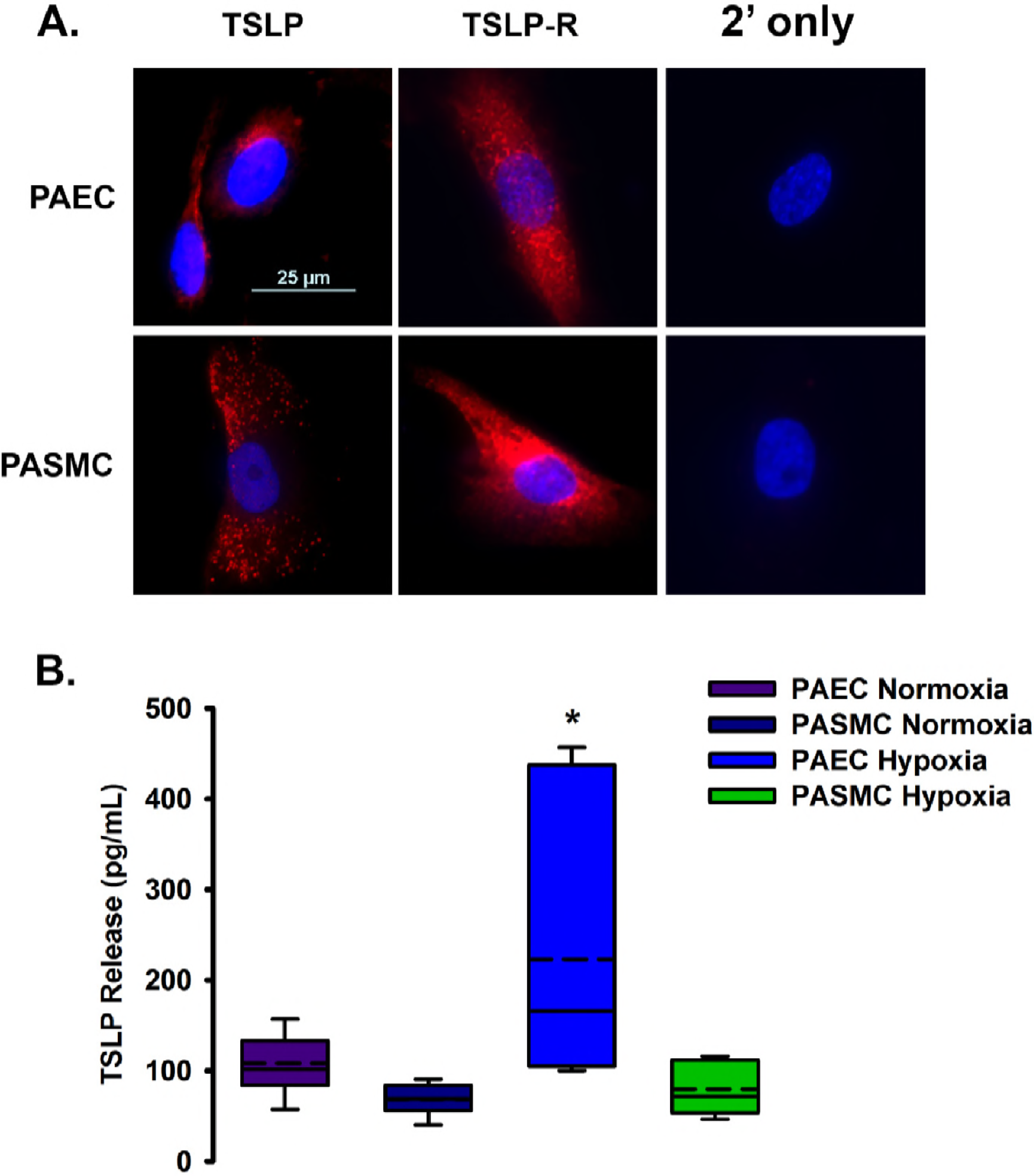
Thymic stromal lymphopoeitin (TSLP) and receptor (TSLP-R) expression and secretion in human pulmonary artery endothelial cells (PAECs) and smooth muscle cells (PASMCs). A) Immunofluorescent staining of PAECs and PASMCs demonstrate the presence of TSLP and TSLP-R. Primary (not shown) and Alexa 555 secondary antibody controls reveal only DAPI staining of cell nuclei and establish antibody specificity. B) Extracellular TSLP was measured by ELISA and shows release of TSLP by both PAECs and PASMCs during normoxia. In PAECs, TSLP release was significantly enhanced following 72h of 1% hypoxia. Hypoxia did not appear to influence PASMC TSLP release. (Values are means ± SE. * indicates significant (p<0.05) effect compared to control, n=5-8)

### TSLP and TSLP-R protein expression in human PAEC and PASMC

Western Blot analysis using anti-human TSLP antibody demonstrated TSLP-R expression in both human PAECs and PASMCs. TSLP protein levels in PAECs were significantly increased (+78%) following hypoxia exposure compared to normoxia controls (p<0.05;n=4). In contrast to PAECs, intracellular expression of TSLP in PASMCs was not significantly changed after 72h exposure to hypoxia (n=4) (Figure 2A). Western analysis from human PAEC and PASMC showed expression of TSLP-R and significant upregulation in both cell types following 72h hypoxia compared to normoxia controls (p≤0.05; n=4) (Figure 2B).

**Figure 2.**
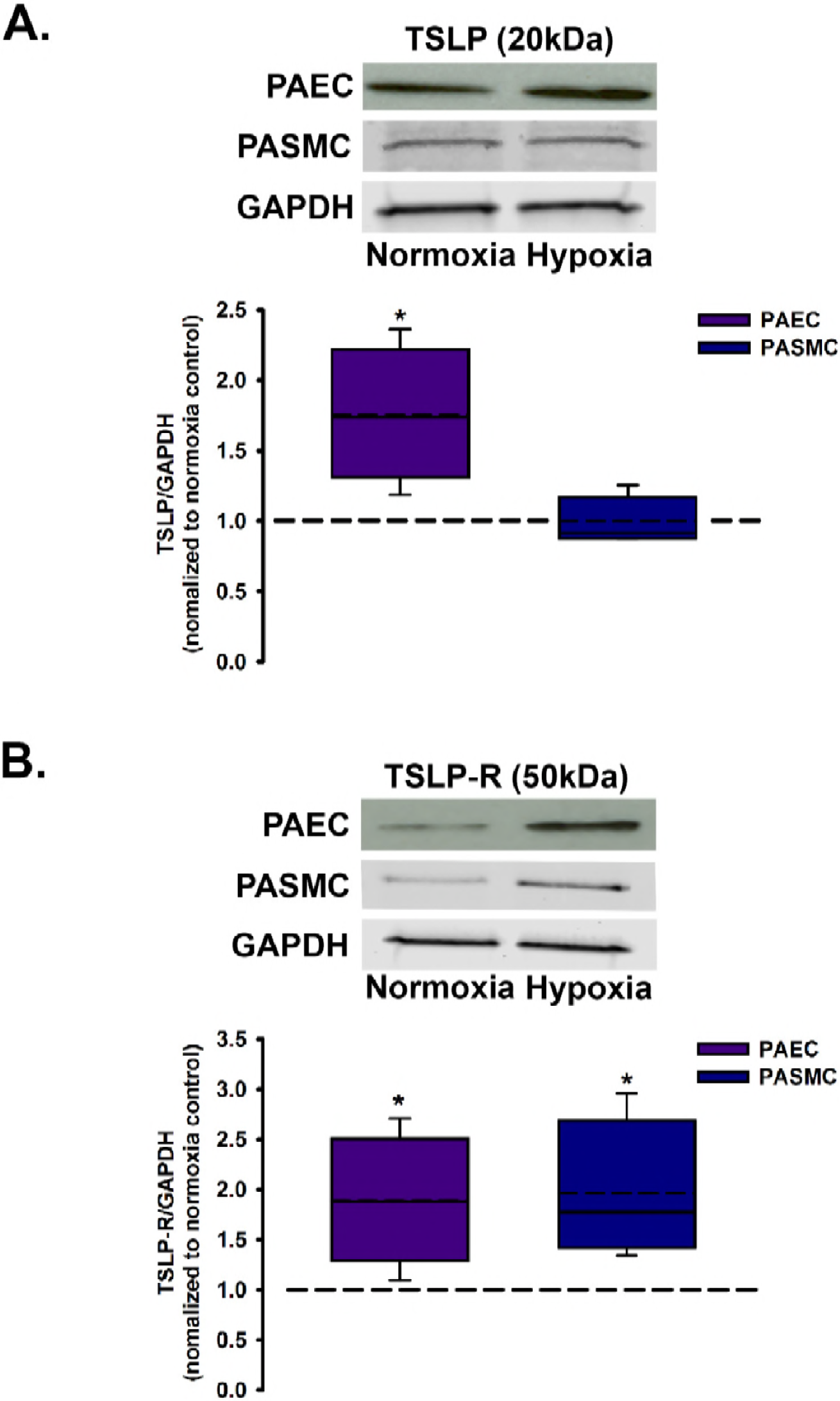
Effect of hypoxia on TSLP and TSLP-R expression in PAECs and PASMCs. A) PAECs exposed to hypoxia for 72 h showed significant upregulation of TSLP expression in comparison to normoxia controls. In contrast, TSLP expression in PASMCs following 72 h hypoxia was unchanged. B) Western analysis showed significant upregulation of TSLP-R expression in PAECs in comparison to normoxia controls. Similarly, TSLP-R expression in PASMCs was increased following hypoxia exposure for 72 h. (Values are means ± SE. * indicates significant (p<0.05) effect compared to normoxia control, n=4 each)

### TSLP enhances human PASMC proliferation

Based on our data suggesting that TSLP, particularly under hypoxia, derives from PAECs, we explored the idea that such TSLP targets PASMCs, and thus focused the remainder of the study on this cell type. PASMCs were seeded into 96 well plates with approximately 10,000 cells/well and proliferation experiments conducted using the CyQuant fluorescent assay. Serum-deprived PASMCs showed baseline proliferation of ~10% over a 72 h period compared to time zero. Human PASMCs exposed to 20ng/ml TSLP showed significantly greater proliferation (+27%) compared to baseline controls. Hypoxia alone also enhanced human PASMC proliferation (+69%), although TSLP in combination with hypoxia had no significant additional effect on proliferation. HIF inhibitor significantly blunted effects on proliferation with hypoxia exposure and in the presence of hypoxia and TSLP (−39% and −54%, respectively). The effectiveness of the HIF inhibitor was confirmed via immunostaining by prevention of HIF-1α translocation to the nucleus (data not shown). Furthermore, pre-treatment of PASMC with LLL12 (STAT3 inhibitor) also inhibited PASMC proliferation with hypoxia and the presence or absence of TSLP (p<0.05, n=4) (Figure 3).

**Figure 3.**
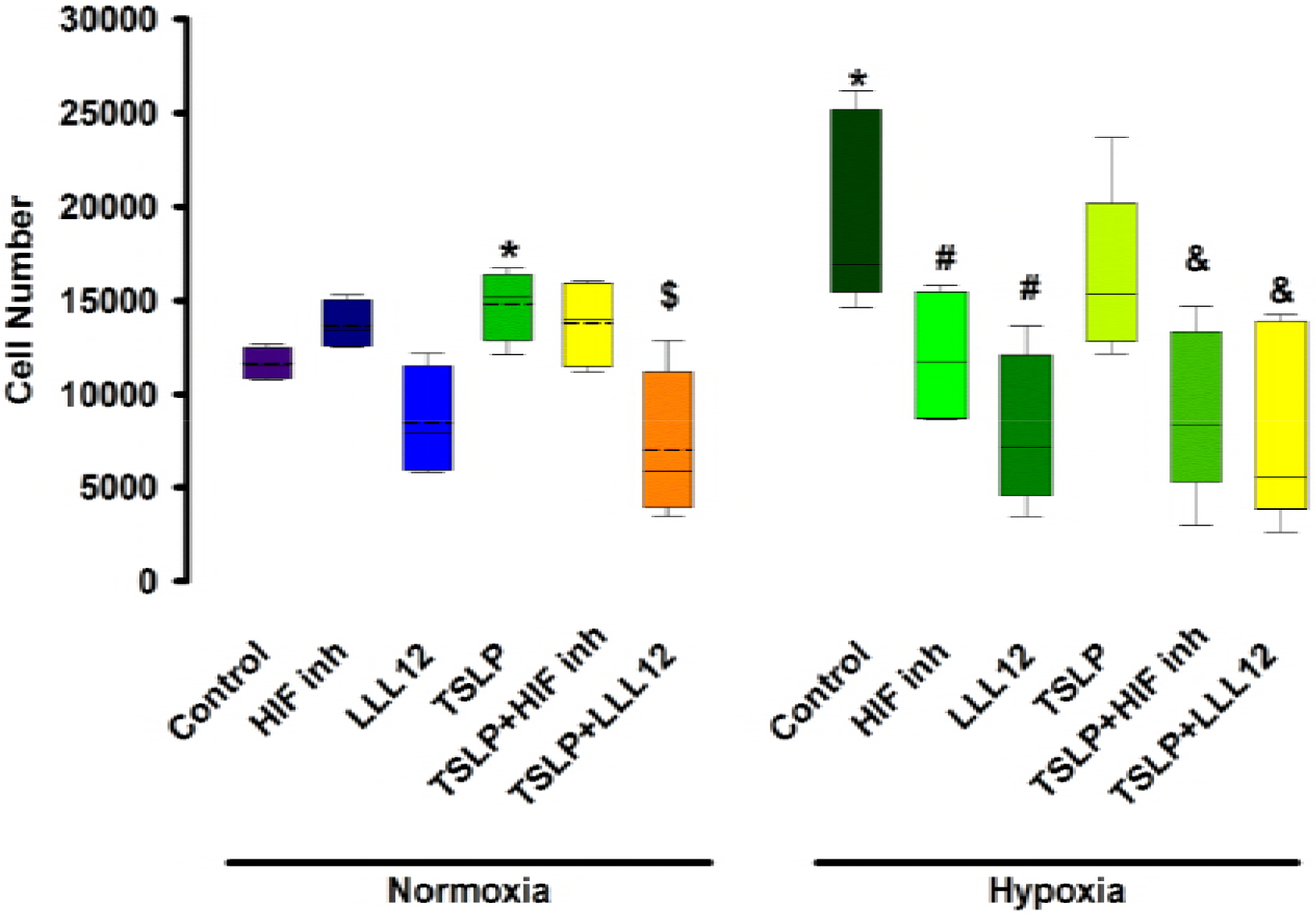
Proliferation of PASMCs. Both hypoxia and TSLP (20 ng/ml; 72h) promote robust cellular proliferation which can be attenuated by inhibition of HIF inhibitor or STAT3 (LLL12) (10μ,M each). See methods for proliferation assay. (Values are means ± SE. * indicates significant(p<0.05) effect compared to normoxia control, # compared to hypoxia control; $ compared to normoxia TSLP treatment; & compared to hypoxia TSLP treatment; n=4 each)

Proliferation was further verified by Western Blot analysis of human PASMCs for proliferating cell nuclear antigen (PCNA) or Cyclin E protein expression following exposure to TSLP or hypoxia, both demonstrating enhanced PCNA (+89% with TSLP and +135% with hypoxia) and Cyclin E (+165% with TSLP and +115% with hypoxia) expression compared to normoxia controls. TSLP had no significant additional effect on expression of PCNA or Cyclin E in the presence of hypoxia. Interestingly, the HIF inhibitor significantly blunted Cyclin E expression in the presence of TSLP during normoxia as well as during hypoxia exposure (−40%). The STAT3 inhibitor, LLL12, significantly reduced Cyclin E expression levels with TSLP treatment in both normoxia and hypoxic conditions (−60% each compared to respective controls). PCNA expression was also significantly inhibited by LLL12 with hypoxia and TSLP treatment (−43%) but was not substantially effected during normoxia (p<0.05, n=5 each) (Figure 4).

**Figure 4.**
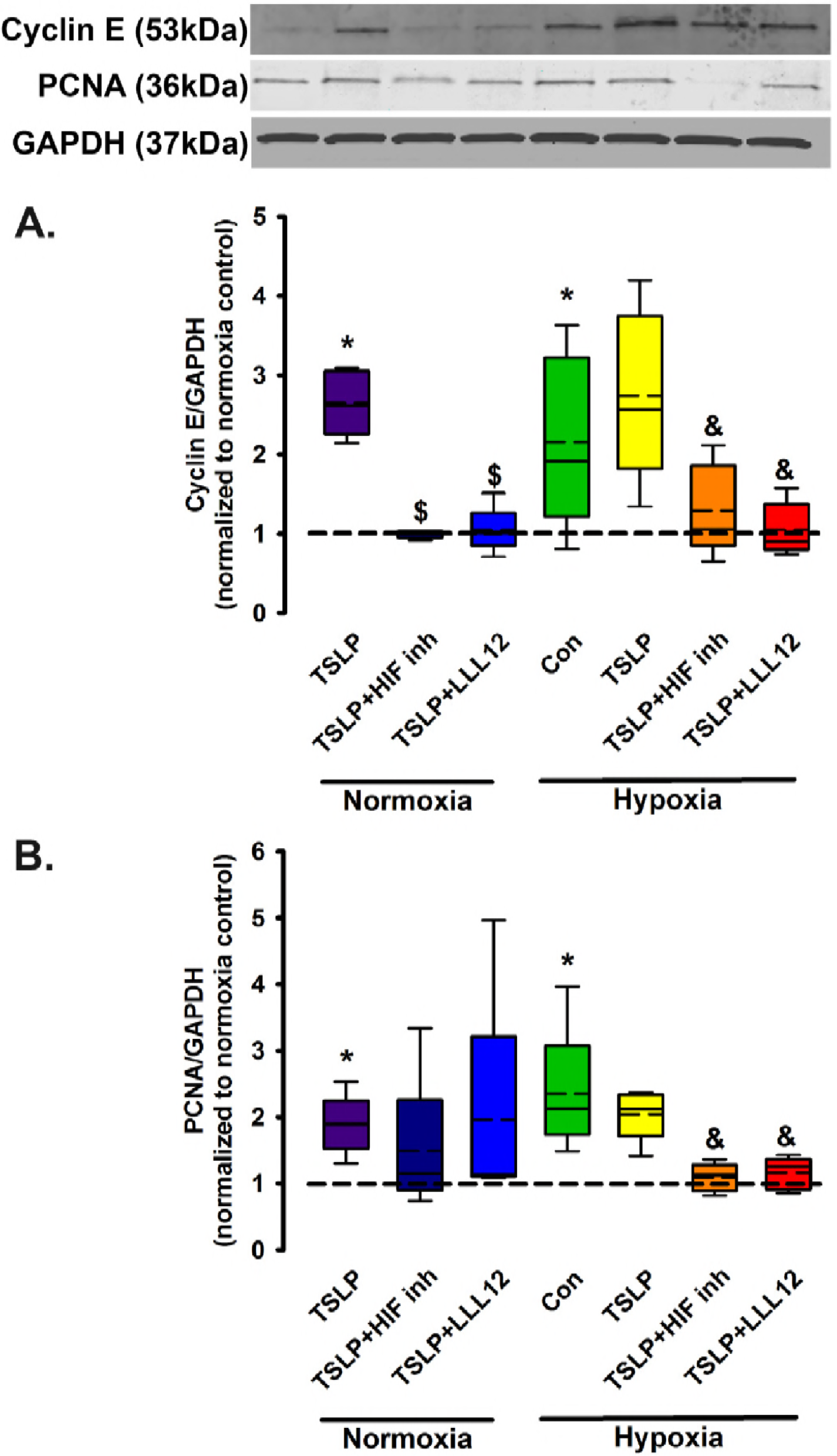
Proliferation of PASMCs in response to hypoxia and/or TSLP. A) Western blot analysis of PASMCs demonstrates upregulation of Cyclin E following TSLP or hypoxia compared to normoxia control. HIF inhibitor and LLL12 in the presence of TSLP significantly blunted increased expression of Cyclin E under normoxia and hypoxia compared to TSLP alone in each condition. B) PCNA expression was also increased with TSLP and hypoxia exposure. HIF inhibitor and LLL12 significantly prevented PCNA upregulation in the presence of TSLP during hypoxia only. (Values are means ± SE. * indicates significant (p<0.05) effect compared to normoxia control, $ compared to normoxia TSLP treatment; & compared to hypoxia TSLP treatment; n=5)

### Mechanisms of TSLP action in human PASMC

MAPK and PI3K/Akt pathways are known to be involved in PA cell proliferation in response to mitogens, thus we investigated the possible link between these pathways with TSLP and hypoxia. Western blot analysis of human PASMCs exposed to normoxia or hypoxia in the presence or absence of 20ng/ml TSLP for 72 h demonstrated no significant change in the expression of ERK1 and ERK2 with hypoxia compared to normoxia. Inhibition of STAT3 significantly blunted ERK1/2 expression (−61% and −64% respectively) under hypoxic conditions and in the presence of TSLP compared to hypoxia+TSLP. Similar trends are observed with LLL12 treatment in normoxia but are not statistically significant. In comparison, PI3K and Akt expression were considerably increased (+125% and +131%) in the presence of TSLP compared to normoxia controls. Additionally, hypoxia also significantly increases expression of PI3K (+240%) and Akt (+228%). HIF inhibition significantly prevented upregulation of Akt (−58%) when exposed to hypoxia and TSLP and blunted PI3K expression when incubated with TSLP in normoxia (−52%) and hypoxia (−68%). STAT3 inhibition with LLL12 substantially reduced PI3K expression (−49%) with TSLP and hypoxia (p<0.05, n=5-6) (Figure 5). Overall, these data suggested several common mechanisms where TSLP and hypoxia intersect.

**Figure 5.**
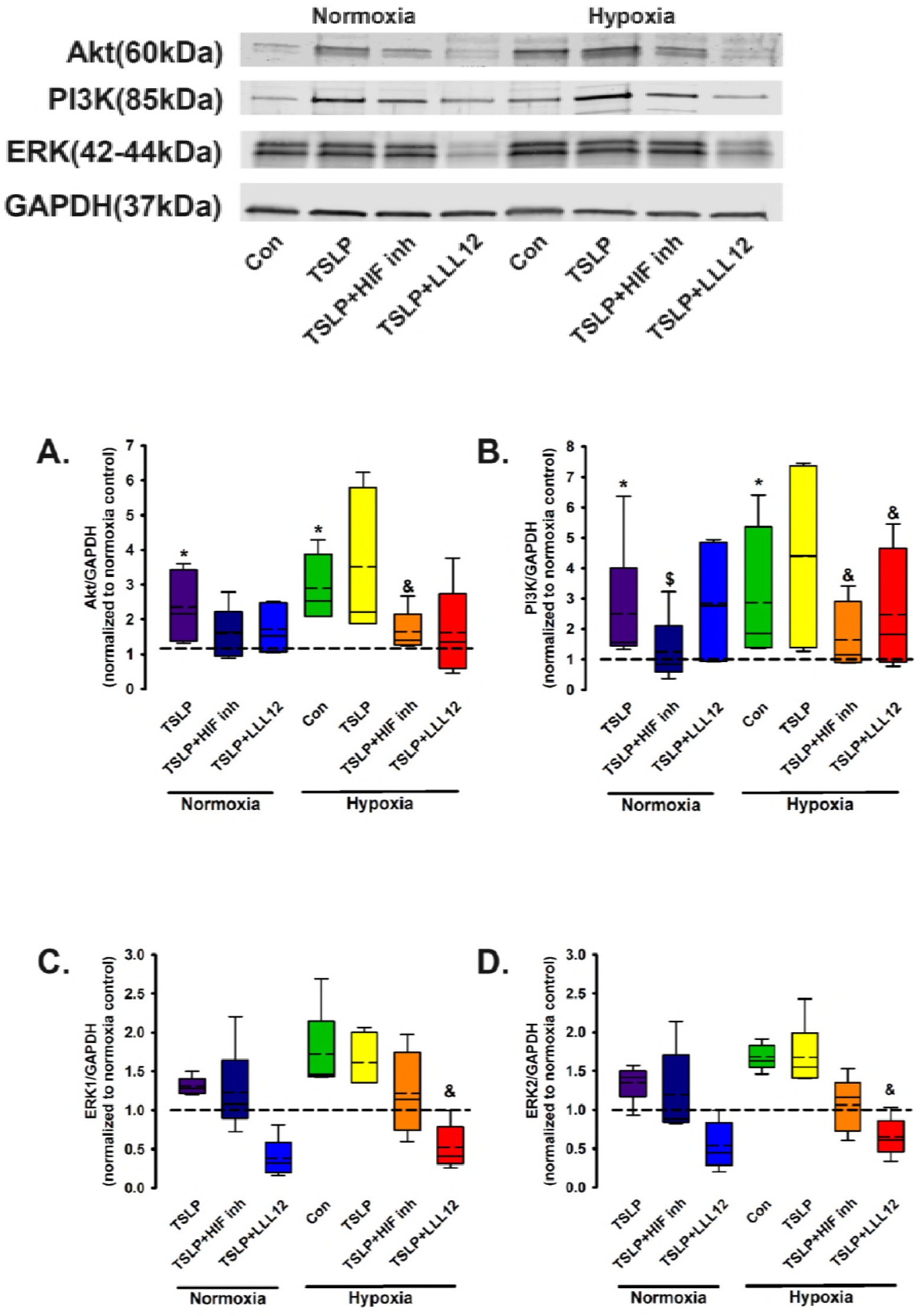
Effects of TSLP on MAPK and PI3K/Akt pathways. A) Western analysis of PASMCs shows increased expression of Akt when exposed to TSLP in normoxia, and hypoxia alone. HIF inhibitor significantly blunted Akt upregulation compared to hypoxia+TSLP treatment. B) Evaluation of PI3K via Western analysis demonstrates significantly increased expression with TSLP in normoxia, and hypoxia alone, an effect blunted by HIF inhibitor compared to TSLP treatment in normoxia and hypoxia. LLL12 prevented PI3K upregulation during hypoxia+TSLP only. C) and D) ERK 1 and ERK2 expression is not significantly altered by hypoxia or in the presence of TSLP in normoxia or hypoxia. LLL12 treatment significantly inhibited ERK1/2 expression in the presence of TSLP during hypoxia. (Values are means ± SE. $ indicates significant (p<0.05) effect compared to normoxia TSLP treatment; & compared to hypoxia TSLP treatment; n=5-6)

### TSLP enhances Jak/STATpathway protein expression in human PASMC

To further delineate the downstream functions of TSLP in human PASMC, we next investigated a known TSLP activated pathway in other cell types: Jak2/STAT3 pathway. Western blot analysis of human PASMCs exposed to normoxia or hypoxia for 72h, with or without TSLP, showed that hypoxia enhances Jak2 expression (+97%) compared to normoxia control. TSLP increased Jak2 expression (+53%) in normoxic conditions but did not potentiate the effects of hypoxia. HIF inhibitor and LLL12 significantly decreased Jak2 expression in the presence of TSLP during normoxia (−38% and −58%, respectively), but only LLL12 treatment caused significant downregulation (−73%) of Jak2 compared to combined TSLP and hypoxia exposure. (p<0.05, n=6) (Figure 6A). In comparison, Western analysis of STAT3 expression shows significant increase with TSLP treatment (+115%) in normoxia and hypoxia (+145%) alone when compared to normoxia controls. HIF inhibitor (−67%) and LLL12 (−70%) significantly decrease the effect of TSLP during hypoxic exposures. (p<0.05, n=6) (Figure 6B)

**Figure 6.**
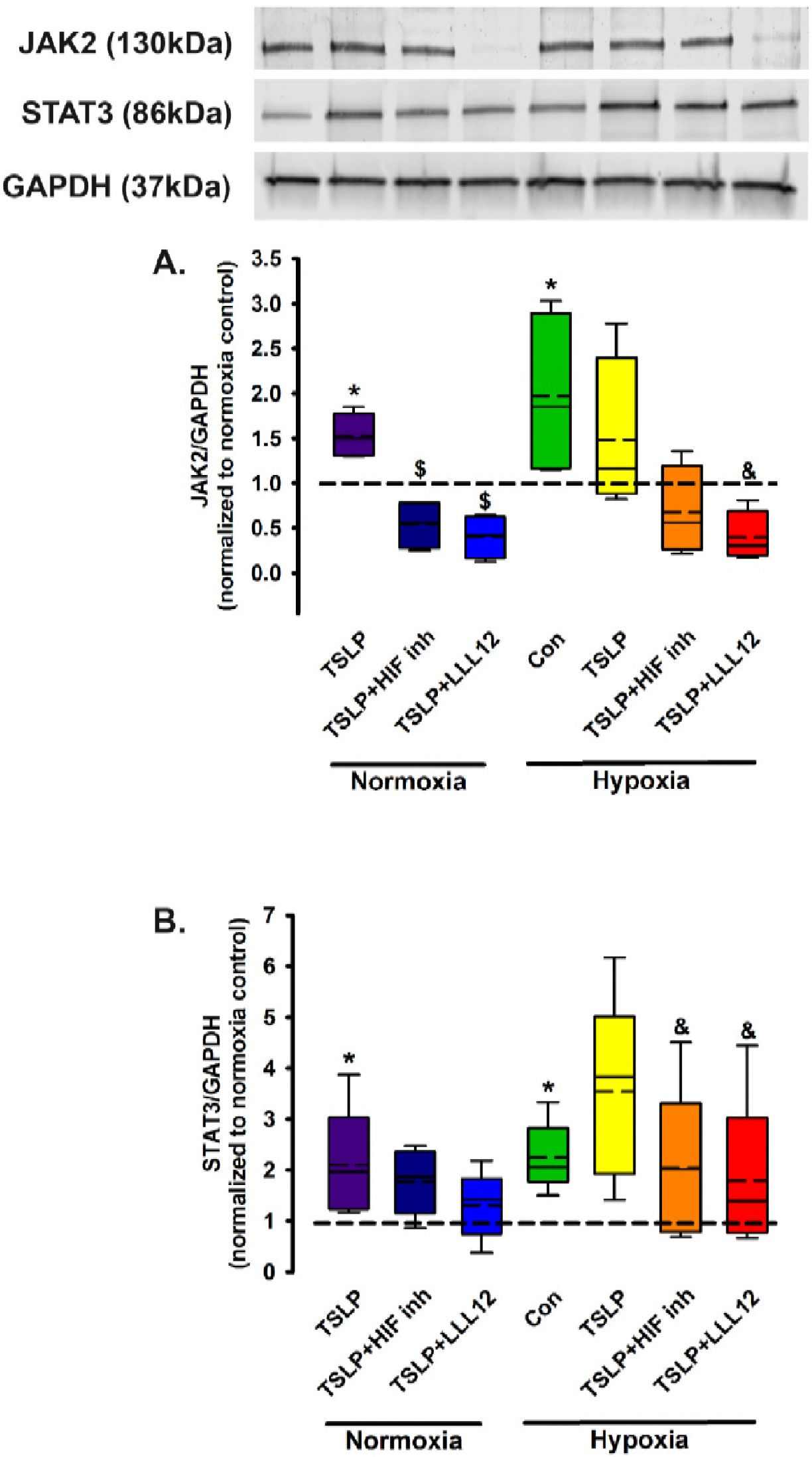
TSLP activates Jak2/STAT3 pathway in PASMCs. A) Western blots of human PASMCs exposed to normoxia, hypoxia, TSLP and inhibitors as previously stated show that hypoxia enhances JAK2 expression compared to normoxia control. TSLP increased JAK2 expression in normoxic conditions but did not potentiate the effects of hypoxia. HIF inhibitor and LLL12 significantly decreased JAK2 expression in the presence of TSLP during normoxia but only LLL12 treatment had significant effects on JAK2 expression compared to TSLP+hypoxia exposure. B) Analysis of STAT3 expression via Western blot show significantly increased expression of STAT3 in normoxic conditions with TSLP and hypoxia alone. LLL12 in the presence of TSLP significantly reduced STAT3 expression compared to TSLP alone. HIF inhibitor and LLL12 substantially reduced STAT3 expression in the presence of TSLP compared to hypoxia+TSLP. (Values are means ± SE. * indicates significant (p<0.05) effect compared to normoxia control, & compared to hypoxia TSLP treatment; n=6)

### Effect of TSLP and hypoxia on [Ca^2+^]_i_ in human PASMC

Enhanced contractility is known to occur in PA during hypoxia or PH; therefore, we examined the role of hypoxia and TSLP on [Ca^2+^]_i_ responses to serotonin in human PASMCs. Exposure to 20ng/ml TSLP for 72 h significantly increased the amplitude of serotonin responses in PASMCs (627±29 nM) compared to normoxia controls (362+15 nM), an effect which was inhibited by HIF inhibitor (493±36 nM). Hypoxia alone significantly increased [Ca^2+^]_i_ responses to serotonin (501±24 nM) which was potentiated in the presence of TSLP (700±41 nM), but blunted by HIF inhibitor (399±38 nM) (p<0.05, n=4) (Figure 7).

**Figure 7.**
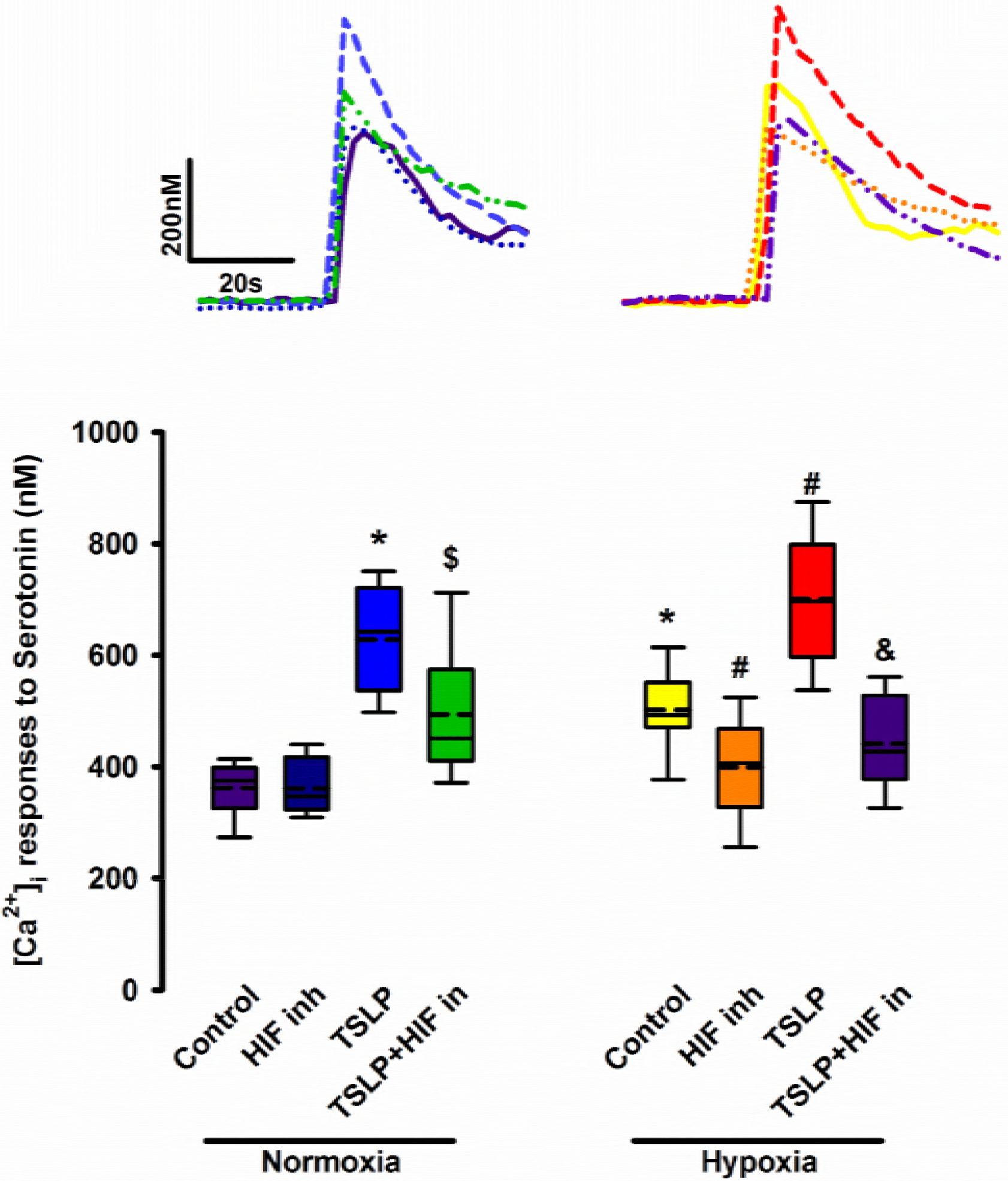
Effect of TSLP on PASMC intracellular calcium ([Ca^2+^]_i_) responses to agonist. Exposure to hypoxia and/or TSLP for 72 h results in increased [Ca^2+^]_i_ responses to serotonin (10μM): effects blunted by HIF inhibitor under both normoxic and hypoxic conditions (Values are means ± SE. * indicates significant (p<0.05) effect compared to normoxia control, # compared to hypoxia control; $ compared to normoxia TSLP treatment; & compared to hypoxia TSLP treatment; n=4)

**Figure 8.**
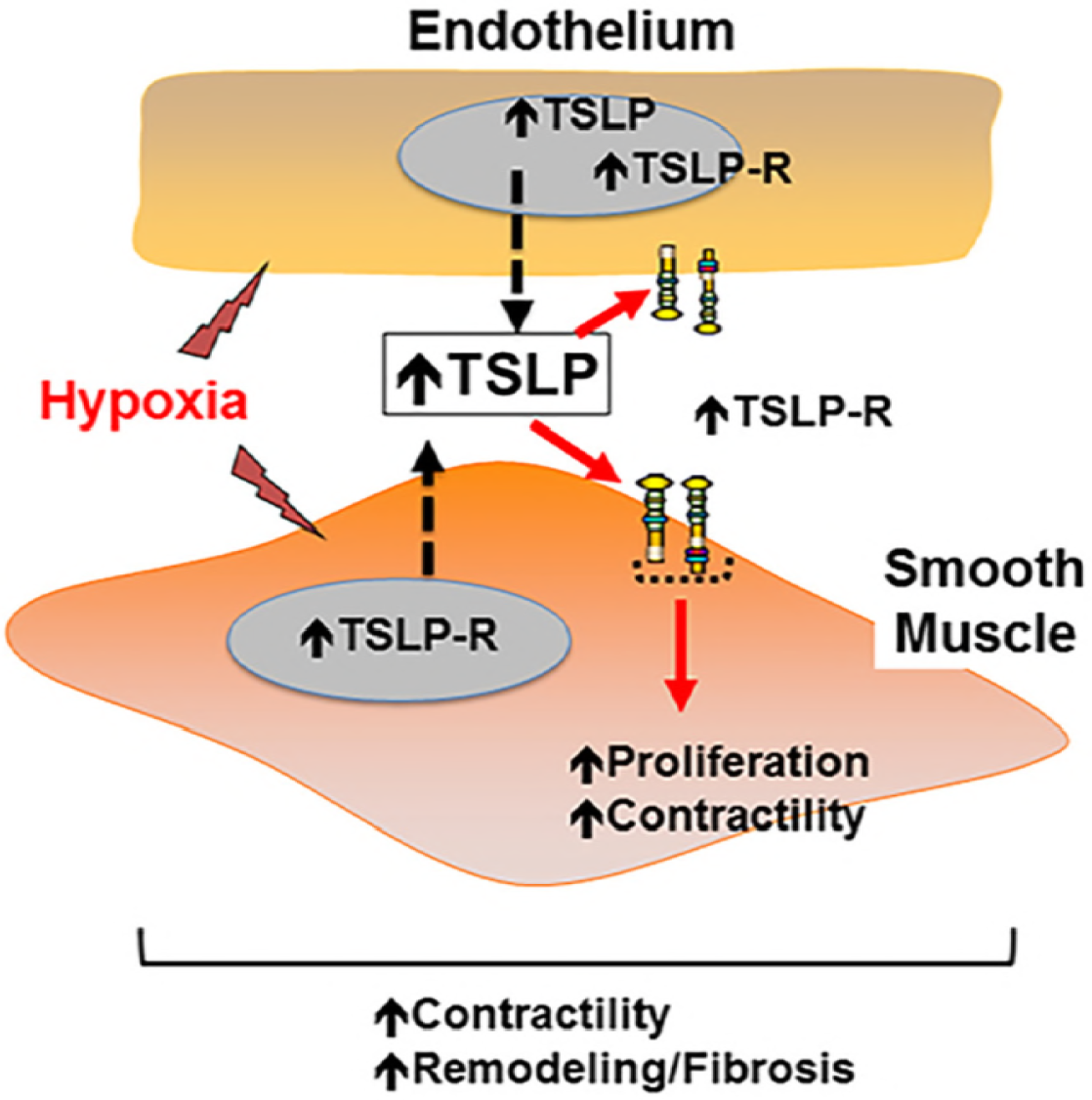
Schematic of TSLP expression and effect in pulmonary artery. TSLP can be derived from PAECs (largely) and PASMCs, with influence on TSLPR expressed by both cell types. Hypoxia enhances PAEC-derived TSLP as well as TSLP-R in both cell types, thus enhancing the potential for TSLP effect. TSLP acts on PASMCs to enhance cell proliferation and calcium responses to agonist (and thus contractility). TSLP may interact with hypoxia by promoting HIF1-α signaling: effects relevant to diseases such as pulmonary hypertension.

## Discussion

In spite of substantial medical advances, PH remains a devastating disease. Although multifactorial in etiology, chronic hypoxia is a well-recognized risk factor and pathophysiologically relevant mechanism in several forms of PH. Furthermore, hypoxia can induce inflammation that only enhances the detrimental structural and functional changes in the PA. Accordingly, understanding the mechanisms that link hypoxia and inflammation, particularly early in disease, is important. In skin and airway disease, there is increasing recognition that TSLP is a locally-produced, critical, early factor that drives the inflammatory cascade relevant to allergy, asthma and chronic obstructive pulmonary disease, and furthermore has non-canonical effects on airway structure and function. Therefore, we hypothesized that locally-produced TSLP mediates and potentiates the effects of hypoxia on PA remodeling and contractility via TSLP receptors present in the PA.

Pulmonary artery endothelial cells regulate both PA structure and function, e.g. via the well-known release of NO towards vasodilation and endothelin towards vasoconstriction and remodeling. The endothelium certainly experiences alterations in oxygen levels and thus plays a major role in mediating and modulating the effects of hypoxia in pathogenesis of diseases such as pulmonary hypertension. Accordingly, understanding endothelial mechanisms that are altered by hypoxia becomes important. While hypoxia can influence the PAEC in several ways, the idea that locally-produced pro-contractile or pro-proliferative factors permits exploration of the concept that such mechanisms can contribute to induction and maintenance of disease. For example, we had previously shown that in response to hypoxia, human PAECs can release the neurotrophin BDNF which can enhance PASMC contractility [31]. In this regard, the effects of hypoxia on endothelial TSLP have interesting parallels. On the other hand, BDNF also enhances endothelial NO [36] and may thus serve a different purpose in the PA. However, similar to our finding of TSLP effects in PASMCs, BDNF has autocrine effects on PAECs [32] where it modulates HIF1α thus priming PAECs in their response to hypoxia. Whether a similar autocrine influence of TSLP occurs in PAECs is not known, and was not examined in the present study. However, the presence of TSLP-R on PAECs may allow for such effects. What remains to be then examined is whether in the context of hypoxia, overall TSLP effects on the PA would result in vasodilation (e.g. via NO) or vasoconstriction due to the effects of TSLP on PASMC calcium responses, as observed in the current study.

Within the PA, smooth muscle cells also play important structural and functional roles through effects on vascular tone and contractility, as well as vascular stiffness and fibrosis via cell proliferation and production of extracellular matrix. Accordingly, mechanisms derived from PAECs as well as PASMCs can have substantial local influences on the smooth muscle in acute and chronic conditions. Here, we demonstrate that TSLP is secreted by both PAECs and PASMCs, although it appears that PAECs are a more substantial source, and importantly it is PAEC-derived TSLP that is upregulated under hypoxic conditions. Thus, we propose that PASMC-derived TSLP may represent a “background” level with perhaps autocrine/paracrine effects. What then becomes important are the effects of TSLP on PASMCs, and our current results show contributions to enhanced PASMC proliferation and contractility, particularly in hypoxia.

The effects of TSLP on PASMCs involve the receptor TSLP-R which is abundantly present in smooth muscle. Interestingly, hypoxia, acting via HIF-1α upregulates TSLP-R, pointing to at least one mechanism by which hypoxia and TSLP could interact, particularly given the additional observation of increased TSLP production by PAECs in hypoxia.

There is currently little information on how hypoxia may regulate TSLP or TSLP-R in any tissue. Data from skin, airways and GI suggest that TSLP regulation is highly cell-type specific (e.g. constitutive vs. induced, responsiveness to specific cytokines such as TNF-α, IL-1) [5, 6, 11]. Even in these systems, much of the information is from cellular mRNA, with substantially less data on TSLP secretion which is required for autocrine/paracrine function. Nonetheless, promoters for HIF-1α have been identified on the TSLP gene,[35, 37, 38] and some studies have shown ERK1/2 mediated alterations in TSLP expression[35, 39] during inflammation. In terms of TSLP-R, there is even less information on its regulation, but a weak HIF-1α promoter is recognized, while the role of other transcription factors is not known. In non-vascular systems TSLP expression can be enhanced in a HIF-1α dependent manner[40] and is consistent with the presence of the promoters. Our studies showing the suppressing effect of HIF-1α inhibitor on TSLP or TSLP-R expression are consistent but further exploration on such regulation is needed. Furthermore, it would be important to determine whether baseline TSLP/TSLP-R expression is different, or differently influenced by hypoxia in patients with PH, especially if mechanisms such as HIF-1α are involved.

PH represents both an imbalance between the extent of vasodilation and vasoconstriction, as well as PA remodeling, and the latter involves PAEC and PASMC proliferation and migration, resulting in dysfunctional endothelial and smooth muscle. *In vitro*, PASMCs and PAECs can proliferate in response to multiple signals. JAKs are involved in such mitogen-induced signaling in human PA [35, 41] Additionally, ERK and PI3K pathways, as well as Rac1, are also important in PA cell proliferation induced by mitogens [35, 42, 43]. These signaling pathways involved in mitogen-induced PA cell proliferation happen to be also involved in TSLP signaling, at least in other non-vascular systems. For example, TSLP induces cell proliferation of the human myeloid leukemia cell line MUTZ-3 via STAT5 phosphorylation [44]. In human airway smooth muscle, we previously showed that STAT5 is activated by TSLP.[5] Similarly, in human airway epithelial cells, MAPK, PI3/Akt and NFκB are all activated by TSLP [45, 46]. The present results showing increase expression of PI3/Akt are entirely consistent in this regard. One report did show that phosphorylation of JAK 1 and 2 precedes STAT3 phosphorylation upon TSLP-TSLP-R binding in human lymphoid cells [44]. Indeed, it is likely that the range of signaling mechanisms activated by TSLP is species-, cell- and perhaps context-dependent. Nonetheless, the role of these signaling intermediates in TSLP effects on PASMC was previously unknown, especially in the presence of hypoxia or underlying PH. What makes TSLP particularly relevant is the increasing interest in STAT inhibitors for a number of diseases such as fibrosis.[47]

An interesting observation in our studies was that even under normoxic conditions, inhibition of HIF-1α resulted in blunting of TSLP effects in PASMCs, for example on intracellular calcium signaling. These data suggest that TSLP may prime PASMCs for hypoxia effects. In this regard, the baseline expression and secretion of TSLP by PASMCs (or PAECs) may play such an autocrine role. While we did not observe a synergistic effect of TSLP and hypoxia on different parameters, we also point out the extremely potent level of hypoxia and the extended duration of exposure in these initial studies. Further exploration using different hypoxia and TSLP exposures are needed to determine the functional significance of TSLP effect on HIF-1α.

In addition to effects on cell proliferation, we observed that TSLP enhances [Ca^2+^]_i_ in PASMCS. Vasoconstriction involves increased [Ca^2+^]_i_ and contractility of PASMCs, which may be mediated by a number of regulatory mechanisms, particularly Ca^2+^ influx mechanisms such as store-operated entry and voltage-gated channels, as well as intracellular Ca^2+^ release from sarcoplasmic reticulum, and enhanced Ca^2+^ sensitivity for force generation, partially involving the RhoA/Rho kinase pathway [48, 49]. Previous studies have already shown the importance of enhanced constrictive but blunted dilatory mechanisms in mediating the effects of hypoxia in the PA, as well as their contribution of PH pathophysiology. Our results suggest a central role for TSLP (e.g. that derived from PAECs) in hypoxia-induced modulation of [Ca^2+^]_i_ and contractility in PASMCs. There is currently little information on such effects of TSLP in any vascular system. However, in recent studies using human airway smooth muscle, we demonstrated that TSLP can enhance [Ca^2+^]_i_ responses to agonist, at least via increased Ca^2+^ influx.[5] Whether TSLP influence RhoA/Rho kinase and Ca^2+^ sensitivity is unknown in PA and will be explored in future studies.

In conclusion, our study demonstrates that local production of TSLP occurs in the PA, largely involving PAECs, while PASMCs which express TSLP-R are responsive. Hypoxia enhances local TSLP production and PASMC TSLP-R expression leading to effects on PASMC proliferation and enhancement of [Ca^2+^]_i_. As both TSLP production and TSLP-R expression are enhanced with hypoxia, the present study suggests that TSLP may play a role in hypoxia induced diseases including PH.

## Sources of Funding

This work was supported a grant from the Mayo Center for Biomedical Discovery (CBD; Prakash).

## Conflict of Interest

None declared.

